# Defining the multiplex probe panel for detecting mutating viruses with high clinical sensitivity

**DOI:** 10.1101/2024.04.23.587389

**Authors:** Hannah N. Kozlowski, Ayokunle A. Lekuti, Muhammad Atif Zahoor, Jordan J. Feld, Warren C. W. Chan

## Abstract

Nucleic acid technology has emerged as an important diagnostic for infectious diseases, cancer, cardiovascular diseases, and other diseases. However, mismatches between the probes and targets can lead to misdiagnosis. Here we determine how many mismatches between the probe and target lead to poor clinical performance and respond by developing a rationale multiplex strategy to overcome this detection problem. We found that the probe-target mismatches of greater than 20% yielded clinical sensitivity of 22% or less, rendering the diagnostic test useless. We designed probe panels to improve the clinical sensitivity. We tested our multiplex probe strategy using hepatitis C virus as the model pathogen because this virus has high mutation rates. We showed that we can improve the clinical sensitivity for detecting hepatitis C virus from 31 to 89% when we designed and applied a four-probe panel to the diagnostic test instead of a single probe system. Interestingly, increasing beyond four probes did not significantly increase the clinical sensitivity. We present a strategy to overcome the poor clinical sensitivity of nucleic acid tests for mutating genetic targets. Incorporating this panel design strategy can lead to improved diagnostic test performance, fewer false negatives and more accurate treatment planning for patients.

## INTRODUCTION

Diagnostics enable the detection of diseases, and the results guide clinicians to properly administer therapeutics to the patient. Simple nucleic acid tests are increasingly accepted as the gold standard tools for diagnosing infections, cancers and other diseases. These tests provide high diagnostic precision because of the uniqueness of a virus or cell’s genetic sequence. In engineering nucleic acid assays, one sequences the cell’s DNA or RNA to create a blueprint for designing the nucleic acid probes. The probes contain a transducing agent plus a capture sequence that recognizes the cell’s genetic sequence (target). In an assay, the probe binds to the cell’s sequence to produce a signal (e.g., optical, electrochemical, or magnetic) that can be measured or visualized. Genetic assays could be in a sandwich format where the target sequence bridges the probe to the secondary detection sequence or direct format where the probe binds to the target to produce a signal. ^1–3^ A signal indicates illness and initiates the treatment process.

There are many different formats for nucleic acid assays, such as microarrays, bead arrays and polymerase chain reaction (PCR). But fundamentally, the probe design is key to ensuring the success of detecting the target with high precision and sensitivity. The analytical performance of nucleic acid assays has been well characterized when there are mismatches between a probe and the target. ^4–8^ Naiser *et al.* showed that a mismatch in the center of a 16 base microarray probe can reduce signal by 60% but this varied between sequences. ^8^ Swanson *et al.* and Chudy *et al.* showed that mutations in the HIV virus allowed the virus to evade detection. ^9,10^ Klungthong *et al.* showed that mismatches between the rapidly mutating influenza virus and PCR probes resulted in false negative results for patients. ^11^ Strategies for bolstering sensitivity include designing multiple probes for multiple viral genes or monitoring genetic drift and updating probe sequences regularly. ^11–13^ These strategies do not incorporate clinical performance, a key measure of diagnostic usefulness, into test design. Upon further analysis, it becomes evident that the impact of probe-targets mismatches on clinical performance is unclear. Understanding the relationship between the binding of the probe with the target sequence will be key to engineering assays with high clinical sensitivity. Specifically, our study answers the question “what is the effect of probe-target mismatch on the clinical sensitivity of an assay?” While our experiments use a microbead assay as a model system, our approach to identifying the clinical performance as it relates to the probe-target binding can be adapted to other nucleic acid assay platforms.

## RESULTS AND DISCUSSION

### The effect of nucleic acid mismatches on detection

We examined the DNA binding between a probe and target sequence to assess the impact of mismatches on target detection. Figure 1a describes our assay. We selected a microbead assay for this study because the microbead provides versatility in analyzing large numbers of nucleic acid sequences and detection is easily determined using flow cytometry.

**Figure 1.**
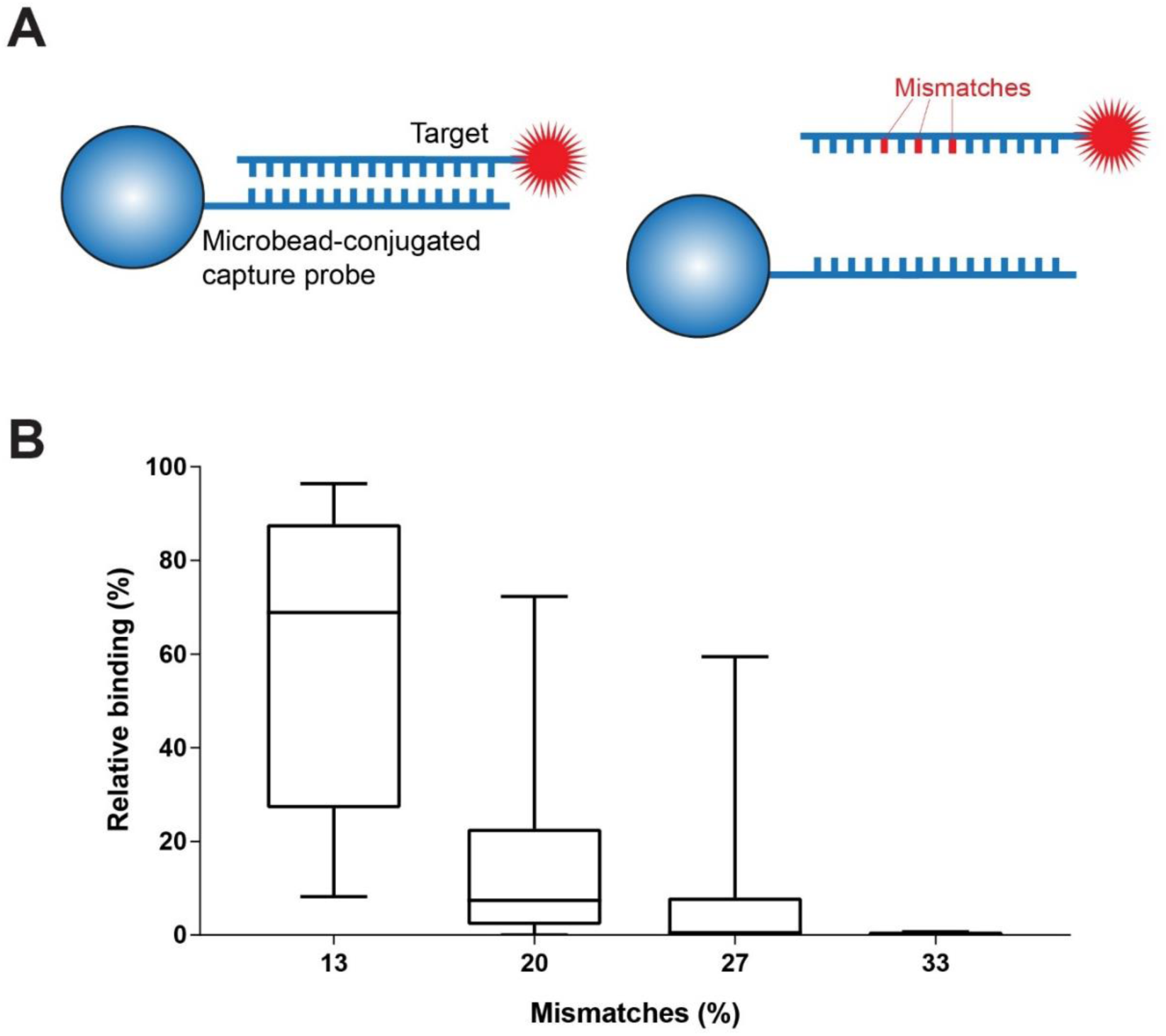
Quantifying the effect of mismatches on binding between microbead-conjugated probes and fluorophore-labelled DNA targets. A) Schematic of the microbead assay. Binding between the bead and fluorophore-labelled target is reduced by mismatches, that can be caused by genetic mutations. B) Box plot depicting relative binding for probe-target pairs with different percentage of mismatches. Measurements were relative to signal from 0% mismatch target. Box plots depict the interquartile range.

We conducted our initial experiments with a nucleic acid sequence of 15 bases (referred to as 15-mer) because diagnostics assays typically use probe lengths between 15-35 nucleotides. ^1,14,15^ We selected the assay temperature at 37⁰C because this temperature is well below the probe-target melting temperature, most clinical laboratories have standard incubators with this temperature, and the reaction would occur faster than at room temperatures.

We designed a series of nucleic acid microbead probes with different numbers of mismatches to target sequences. The 2.8µm bead was conjugated with a capture sequence (referred to as a probe) and then mixed with a solution of fluorescently labelled target sequence (referred to as a target). The binding of the target to the probe leads to a change in the fluorescence signal of the microbead. The overall fluorescence signal of the microbead and thereby the amount of target bound to the probe surface was measured using flow cytometry. We show that the amount of target per microbead decreased when there was an increase in the number of mismatches (Figure 1b). This result occurs because there are fewer nucleotide bonds between the target and capture sequences. The median relative binding for targets with 13%, 20%, 27%, and 33% mismatches was 61%, 7%, 2% and 0%, respectively. Equation 1 shows how we calculated relative binding. The relationship between percent mismatches and binding was modeled using non-linear regression (**Figure S2a**). The goodness of fit for the model was calculated using standard error of the regression (S=22.1%). We presented figure 1b as a range where the majority of the datapoints are in the boxes. The lines above represent the range of the data points and are not error bars. We expected a large spread in DNA binding because of changes in DNA secondary structures, percent guanine-cytosine (%GC) content and mismatch locations due to the probe-target mismatches. When looking at the median results (in the boxes), there was a trend toward decreasing relative binding with increased mutations. We observed the greatest decrease in binding when the targets contained 20% mismatches to the probe sequence. When targets contain ≥20% mismatches, the results of a nucleic acid assay may not be reliable because there is less than 10% binding to the probe.

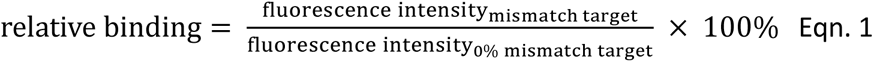

We further explored the effect of the positions of mismatches on binding. We calculated the number of complementary base pairs between mismatches and the position of each mismatch in the target sequence. We saw that the amount of target bound to a probe may be greater for adjacent mismatches versus mismatches separated by one or two base pairs (Figure S1a). The difference in the mean binding for each group was not statistically significant (p=0.12). When we calculated the average mismatch position for targets containing more than one mismatch, we found that mismatches near the ends of the target have little effect on binding (Figure S1b). Mismatches in positions 6-12 are close to the center of the target sequence. These mismatches can decrease probe-target binding by more than 60%. This trend helps to explain why targets with the same number of mismatches have different binding affinities to a probe. We modeled the relationship between mismatch position and binding using a linear regression and saw a weak correlation (S=28.6%). The effects of mismatch position and spacing on binding are small compared to the effect of mismatch quantity (Figure S2). Thus, we focus our studies on the effects of mismatch quantity.

Next, we examined the relationship between relative binding and mismatch quantity using 20-mer and 34-mer sequences. We did not use sequences longer than 35 nucleotides because many nucleic acid assays do not use longer sequences due to increased potential for secondary structures and issues with binding kinetics. We conjugated 27 different capture sequences to microbeads to generate 27 unique probes. Capture sequences were designed to be complementary to one target and contain multiple mismatches relative to the other 26 targets. We mixed each probe with each of the 27 different target sequences in a reaction vial. As a result, we assessed 729 (27 × 27) possible probe-target pairing (Figure S3a). Figure 2 shows the effect of mismatch quantity on binding using 20-mer probes. The median binding for targets with 5%, 10%, 15% and 20% mismatches was 76%, 35%, 18% and 3% respectively. We saw probe-target binding decrease as the number of mismatches increased. The greatest decrease in binding occurred in targets containing 20% mismatches. We similarly examined binding using 34-mer probes and targets with mismatches. Figure S3b shows that binding in 34-mers decreased as the number of mismatches increased. We checked whether other solid-phase DNA binding technologies could bind targets with >20% mismatches. We saw that >20% mismatches are poorly tolerated in microarrays and when using bead-based enrichment prior to sequencing. ^16,17^ Using our set of 20-mer and 34-mer probe-target pairs we saw that a single probe can tolerate <20% mismatches.

**Figure 2.**
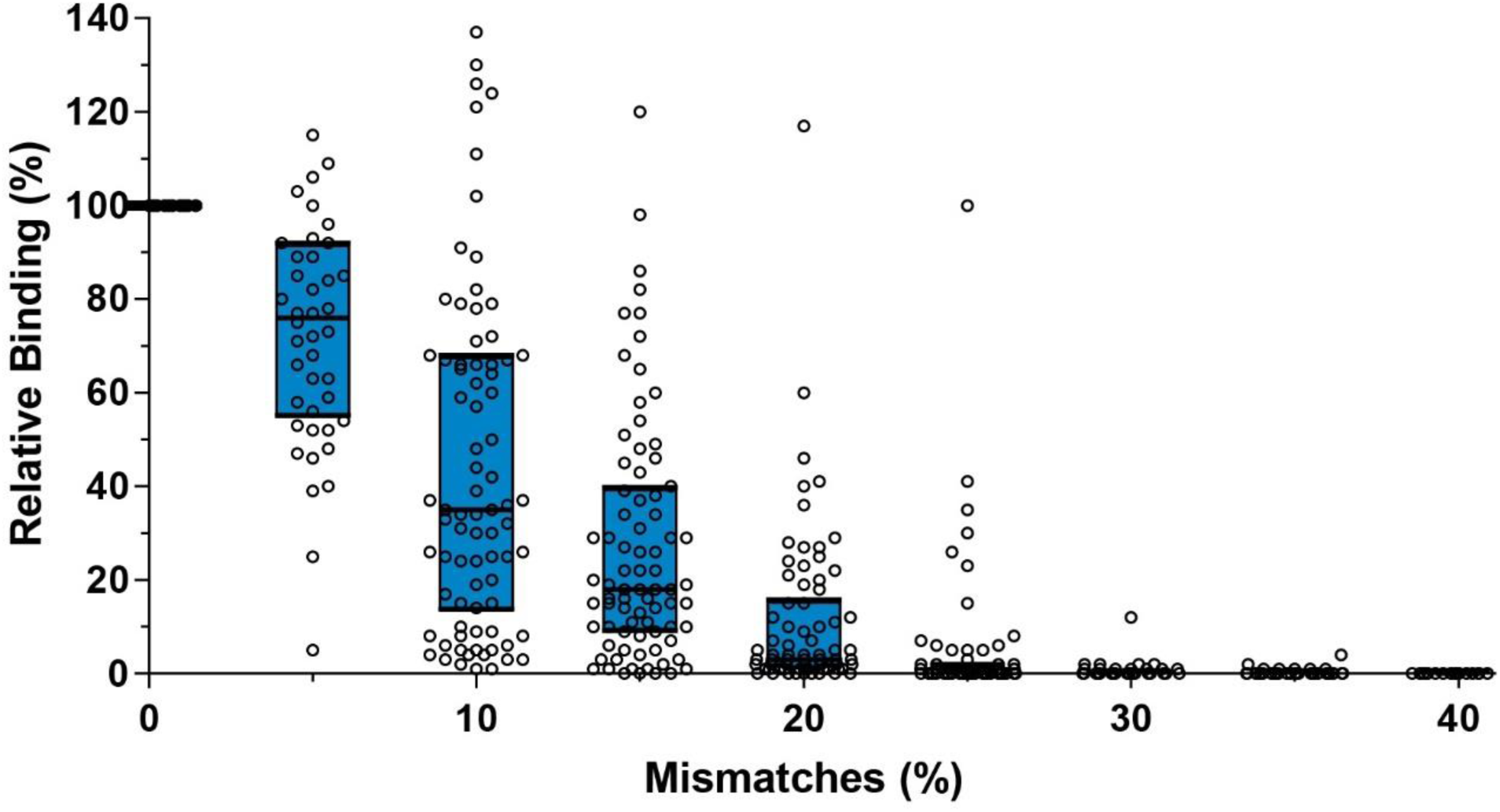
The relationship between mismatch quantity and detection using probe-target pairs with a 20-nucleotide binding region. Relative measurements of probe-target binding for probe-target pairs with increasing percentage of mismatches. Bars represent the 25^th^, 50^th^ and 75^th^ quartiles. Targets with <140% relative binding are shown.

We explored the effect of target concentration on the relationship between relative binding and mismatch quantity using the 20-mer probes. We tested different concentrations of target while holding the number of probes per reaction constant. This simulated the range of pathogen loads seen between patients or during different phases of infection.^18^ We tested 5nM, 15nM, 150nM and 400nM of target which spanned the dynamic range of the microbead assay. As the number of mismatches increased probe-target binding decreased irrespective of concentration (**Figure S4**). Targets with concentrations within the linear range and containing 20% mismatches obtained 1% to 19% relative binding. Targets with 20% mismatches and concentrations above the linear range obtained 21-131% binding relative to 15nM perfectly complementary targets. Increasing the target concentration above the linear range (to 150nM or to 400nM) could increase relative binding for 20% mismatched target but not for targets with greater than 20% mismatches. We saw that targets with >20% mismatches were poorly detected irrespective of concentration.

### The impact of mismatches on clinical sensitivity when using a single capture probe

We wanted to determine the impact of mismatches between the probe and target on the clinical performance of an assay. We aimed to determine the relationship between viral mutation and clinical sensitivity. Clinical sensitivity is a measure of diagnostic performance that indicates the proportion of positive samples that are correctly identified as positive relative to the reference test (equation 2). ^19^ Clinical specificity was not evaluated because probe sequences were designed to bind genetically diverse regions of viral genomes, instead of conserved regions that can be similar between viruses. We selected a rapidly mutating virus as a model system for this study because it provides us with a greater range of nucleic acid sequence variability. We designed 27 unique probes (labelled C1-C27) that were complementary to the core region (genomic region encoding the core antigen) of the hepatitis C virus from 64 infected patients. Each sequence had a guanine/cytosine content of 57-74% and 0-13 mismatches to a probe. We incubated the probes with the target for 30 minutes and measured the microbead’s fluorescence signal via flow cytometry. We analyzed our samples to calculate the clinical sensitivity. As expected, the clinical sensitivity was 100% when the target and probe sequences were 100% complementary to each other (Figure 3a). The median clinical sensitivity for targets with 5% and 15% mismatches was 100% and 89%, respectively. The clinical sensitivities declined as the number of mismatches between probes and targets increased. The median clinical sensitivity for samples with 20% mutations was only 22%, while the clinical sensitivity for samples with >30% mutations was 0%. The presence of highly mutated targets reduced the clinical sensitivity of a diagnostic test. We conclude that a 20% mismatch between the probe and target would lead to clinical performance below the acceptable minimum for a molecular diagnostic test. ^20–22^

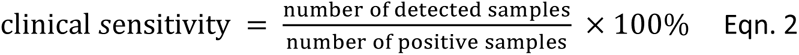

**Figure 3.**
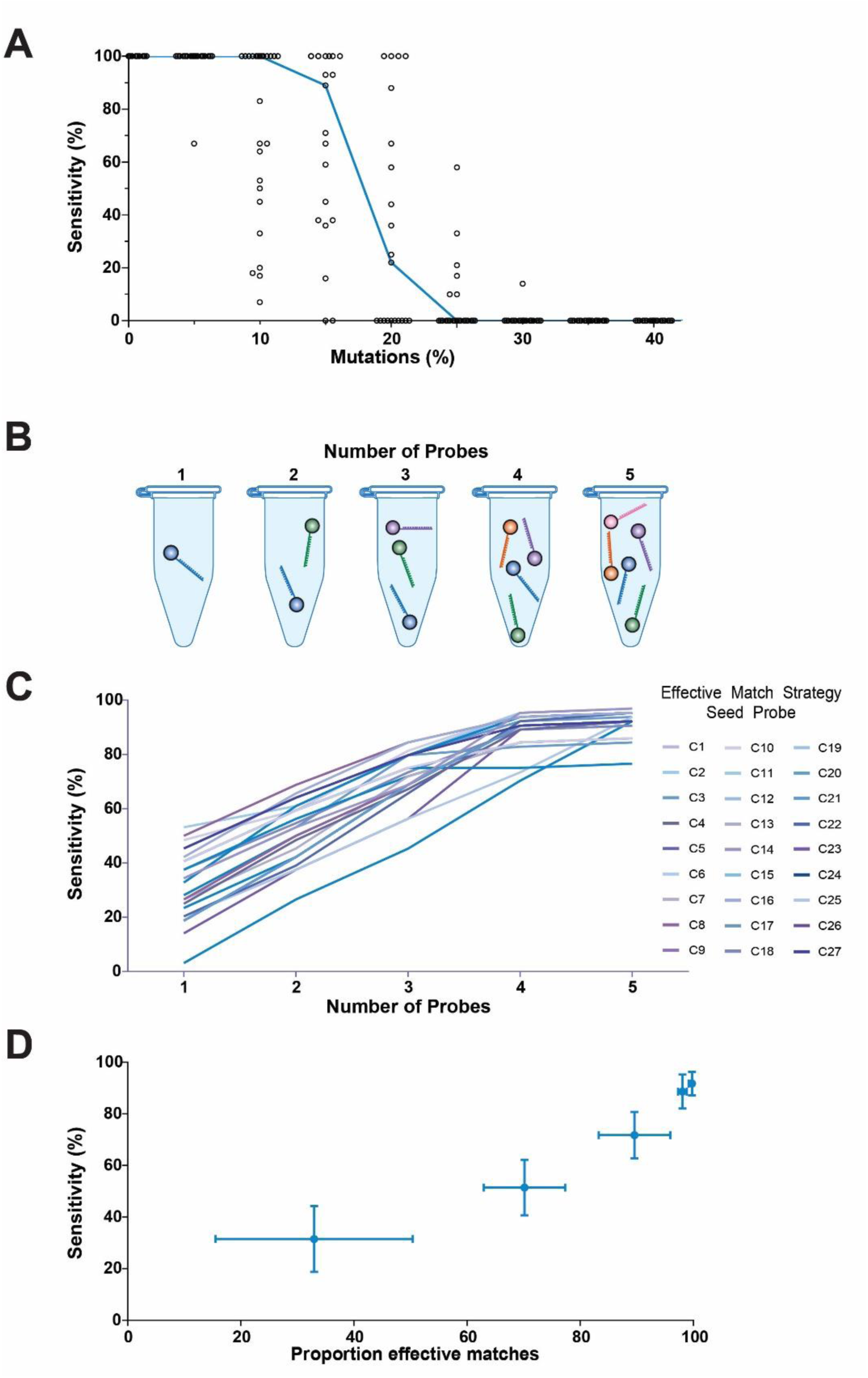
Maximizing clinical sensitivity for detecting a highly mutated region of the Hepatitis C virus core gene. A) Sensitivity versus percent mutations are depicted for probe-target pairs containing ≤40% mutations. B) Schematic of multiple probe assays. C) Line graph showing the sensitivity for the multi-probe combinations. Each line appears in a different color and represents different iterations of the effective match strategy. Each iteration uses a different seed (starting) probe. D) Proportion of effective matches versus sensitivity for 1, 2, 3, 4 and 5 probe combinations. Probe combinations were identified using the effective match strategy. Probes contain between 0-65% mismatches with targets depending on the probe-target pair. Error bars show standard deviation.

### A strategy to mitigate the effect of mutations on clinical sensitivity

A central question then is what strategies can increase clinical sensitivity when probes are greater than 20% mismatched to the target sequence. We hypothesize that the use of multiple probes will lead to large improvements in the clinical sensitivity when there is a 20% or greater mismatch between the probe sequence and target sequence. We tested the impact of strategic multiplexing (the detection of multiple targets simultaneously) on the clinical sensitivity of the microbead assay using probes for the core region. For probe-target mismatches of ≥20%, we asked if the use of multiple probes would alleviate the poor clinical sensitivity for the core targets. We developed a strategy (referred to as the effective match strategy) for building high sensitivity multiplex panels from single probe assays by sequentially adding probes. We began by quantifying the number of mismatches between the initial probe and each target. Targets with fewer than 20% mismatches to the probe have a high probability of detection and are called *effective matches*. We then calculate the number of effective and non-effective matches to that initial probe. The non-effective matches are unlikely to be detected by the initial probe so we design a library of probes capable of binding the non-effective match targets. We repeat the process of identifying mismatches and quantifying the number of effective matches for the candidate probes in the library. We quantify the number of *effective matches* for each candidate and the candidate probe that maximally increases the effective matches is chosen to be the 2^nd^ probe in the multiplex (additional description in Methods). The process is repeated to obtain 3^rd^, 4^th^, 5^th^, *etc*. probes. Unlike other strategies for creating multiple probe assays, our strategy is based on the relationship between clinical sensitivity and mismatch quantity. ^23,24^ The effective match strategy is used to create high sensitivity nucleic acid assays by combining probes that maximally increase the number of probe-target pairs with <20% mismatches.

We applied this effective match strategy to create two-to five-probe multiplex panels using probes C1 to C27. We first calculated the clinical sensitivity of the core probes before the effective match strategy was applied. The mean clinical sensitivity for a single probe microbead assay was 31% (Figure S5). Next, we calculated the number of detected samples when two probes were combined. The mean clinical sensitivity increased from 31% to 51%. We repeated the effective match strategy and saw a gradual increase in clinical sensitivity (Figure 3bc, S5). We added third, fourth and fifth probes and clinical sensitivities increased by an average of 21%, 17% and 3%, respectively (Figure 3d). Clinical sensitivity was significantly different when using different numbers of probes (p<0.0001). The addition of second, third and fourth probes increased clinical sensitivity (p<0.0001) but the clinical sensitivity of a five-probe versus a four-probe panel was not significantly different (p=0.72). Increasing the number of probes in a multiplex assay increases complexity, increases optimization time and can increase the number of false positives. An ideal multiplex assay uses few probes to achieve a clinically relevant sensitivity and low cross-reactivity. Here we showed that increasing the number of probes does not inherently increase clinical sensitivity. Multiplexing increases clinical sensitivity but the increase in clinical sensitivity becomes less with each additional probe.

### Validating that the probe combination strategy improves clinical sensitivity when detecting mutating viruses

We sought to validate whether a single probe could detect targets with <20% mismatches using structurally different DNA probes. We designed 28 capture sequences (labelled N1-N28) for the nonstructural protein 5B (NS5B) region in hepatitis C virus. Capture sequences for the NS5B region were designed to have less negative self-dimer stability and higher hairpin melting temperatures than the core probes (Figure S6). We measured binding between DNA-conjugated microbead probes and NS5B targets and observed a decrease in binding as the number of mismatches increased from 0% to 55% (Figure 4a). The median binding between probes and targets with 0%, 10%, 20% and 30% mismatches was 100%, 66%, 25% and 0%, respectively. We plotted the clinical sensitivity versus target mismatches for each NS5B probe in figure 4b. The clinical sensitivity was 100% for targets without mismatches. Clinical sensitivity dropped as targets contained more mismatches. When detecting targets with 20% mismatches, several probes had low or 0% clinical sensitivity. Clinical sensitivity for targets containing ≥35% mismatches was 0%. Similar to the core region, we conclude that a single probe for the NS5B region could not reliably detect targets with ≥20% mutations.

**Figure 4.**
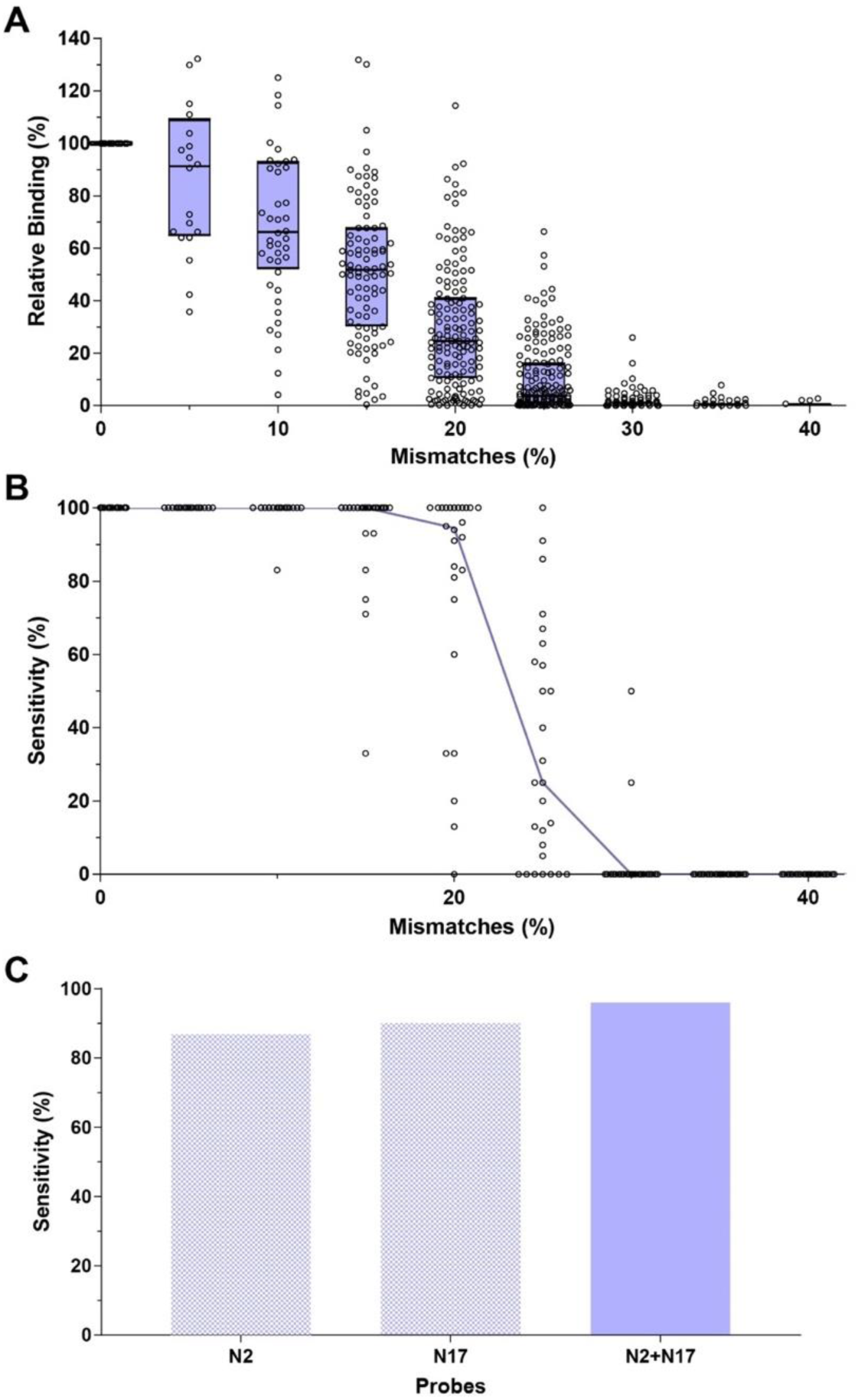
Validating effects of mutations on binding and clinical sensitivity using probes for the NS5B region. A) Relative binding for probe-target pairs with increasing numbers of mismatches. Bars represent the 25^th^, 50^th^ and 75^th^ quartiles. Measurements were relative to signal from the same probe with its perfectly complementary target. B) Sensitivity of each probe for the NS5B region when detecting targets with the given number of mismatches. C) The clinical sensitivities of the N2 and N17 quantum dot barcodes when detecting PCR amplified samples. The purple and blue bars represent single probe versus two-probe detection, respectively.

We designed a proof-of-concept experiment to confirm that the effective match strategy could be used to create a multiplex panel using quantum dot barcodes to address the problem of poor clinical sensitivity for targets containing ≥20% mismatches. Quantum dot barcodes are polymeric microbeads infused with different emitting quantum dots. ^1,25^ Each barcode is conjugated with a different capture sequence. The emission color identifies the capture sequence, which indicates target identity. The first barcode in our multiplex was conjugated with the N2 capture sequence. The N2 sequence had 0-40% mismatches with sequences in our initial group of patients. We applied the effective match strategy to identify a second probe for the multiplex. We conjugated a second barcode with the N17 capture sequence. Instead of using a secondary (reporter) probe to create a sandwich assay, we created a simpler quantum dot barcode assay. DNA targets were labelled during amplification using a fluorophore-conjugated forward primer. We collected samples from a second group of patients to test the ability of this barcode combination to detect hepatitis C. Patients were infected with hepatitis C genotypes 1a, 1b, 2 or 3 and had viral loads from 20 IU/mL to 10^8^ IU/mL. Hepatitis C virus was extracted, reverse transcribed and PCR amplified. We detected the labelled hepatitis C amplicons using the quantum dot barcode assay. The clinical sensitivities of the N2 and N17 barcodes were 91% and 87%, respectively (Figure 4c). We combined the N2 and N17 barcodes to obtain 96% clinical sensitivity. We tested clinical specificity using hepatitis B virus and SARS-CoV-2. The N2 and N17 probes had 100% and 91% specificity, respectively (Table S3). By using the effective match strategy, we decreased the number of targets with ≥20% mutations and showed that the effective match strategy can be used to increase clinical sensitivity when detecting clinical samples.

## CONCLUSION

We determined the impact of probe-target mismatches on the clinical performance of microbeads using hepatitis C as a model disease target. Our results show that a ≥20% mismatch between the probe and target will lead to clinical sensitivities below the recommended minimum sensitivity.^21^ The clinical sensitivity decreased from 100% for perfectly complementary probe-target sequences to ≤22% for targets with ≥20% mismatches. Most pathogens can mutate, and different pathogens will mutate at different rates. ^26,27^ These mutations can lead to a high degree of non-complementarity between a probe and target. In effect, a nucleic acid assay is less sensitive as the number of probe-target pairs with high numbers of mismatches increases, which would increase the number of false negatives and lower the clinical sensitivity. Using capture sequences of varying lengths, we show that the diagnostic sensitivity decreases proportionately to the number of nucleotide mismatches. However, we show that the clinical sensitivity can be rescued by using the effective match strategy. This strategy leverages probes that have less than 20% mismatches. The number of probes is dependent on the degree of mismatch between the target and probe. This study suggests that disease targets with large numbers of mutations will require a careful multiplexing scheme to achieve high clinical sensitivity.

Designing and using multiplex bead panels can be simple. Once the bead probes are engineered, the different beads are placed in a reaction tube and patient sample can be added to the tube. We can further simplify the bead panel by placing the beads and buffers into a pre-measured tablet. A clinician or technician would just need to add the sample. The novelty of the current study is that it establishes a relationship between probe-target mismatch and clinical sensitivity. The results will guide the design of multiplexing panels for detecting highly mutated targets. Our findings advance the development of multi-probe systems for diagnosing disease because it allows us to determine the degree of multiplexing (*i.e.*, number of barcodes) required to detect targets that are highly mutated. Further studies would need to investigate different assay conditions and sequences to adapt the effective match strategy to other nucleic acid assays and create a more generalized rule. However, our study presents a framework by which these studies could be done. Finally, while we used quantum dot barcodes as a model multiplexing detection system, there are many other emerging multiplex tests with similar hybridization conditions (surface enhanced raman spectroscopy, NanoString, CRISPR diagnostics) that can also detect mutating biological targets.

## METHODS

### Probe design

Microbead probes were built using amine-terminated single stranded DNA and carboxy-terminated microbeads. DNA probes contained a 5’ amine group attached to a C12 spacer, a 10 nucleotide polyT spacer and a 15, 20 or 34 nucleotide binding region. DNA probes and their complementary fluorescein-labelled targets were stored in 1× TE (pH 8) (Integrated DNA technologies, Coralville, USA). A list of probes for the core and NS5B regions can be found in Table S1. Amine-terminated DNA was conjugated to 2.8µm carboxy-terminated microbeads (Invitrogen, Waltham, USA) or 4.8µm quantum dot barcodes using carbodiimide chemistry. ^28^ DNA was conjugated to 2.8µm microbeads using a modified manufacturer’s protocol. 10µL of microbeads were transferred to a new tube and washed twice with 100µL 25mM MES (pH 5). Supernatant was removed and 2.2µL 100µM capture DNA and 9µL 25mM MES (pH 5) was added to the washed beads. The tube was mixed and incubated with a slow rotation at room temperature for 30 minutes. Immediately before use EDC was dissolved in 100mM MES (pH 5) to a concentration of 50mg/mL. 8µL of EDC solution was added to the microbeads. The tube was vortexed and incubated for 2 hours at room temperature with a slow rotation. Microbeads were washed using 50mM TRIS (pH 7.4) with 0.1% Tween-20 three times. Beads were resuspended in TRIS and stored at 4⁰C. Quantum dot barcodes were synthesized via concentration-controlled flow focusing and conjugated using one-step EDC chemistry. ^28^ Briefly, a solution of 300mg/mL of EDC in 100mM MES (pH 5.5) was prepared. 10^6^ quantum dot barcodes, 2.4µL 10µM capture DNA and 32µL EDC solution were mixed. 100mM MES (pH 5.5) was added until the volume reached 100µL. Solutions were vortexed then incubated for 4 hours at room temperature with a slow rotation. Quantum dot barcodes were washed twice using 500µL wash buffer (0.5× SSC, 0.1% SDS), resuspended in PBS-T (1× PBS, 0.05% Tween-20) and stored at 4⁰C.

### Microbead probe-target hybridization assay

Probes and targets were hybridized in accordance with established protocols. ^1^ Targets were fluorescein-labelled single stranded oligos in 1× TE and the negative control was 1× TE. All targets in a probe set were diluted to the same concentration. The upper bound of the linear range was identified for complementary targets and all targets were diluted to 20nM, 15nM or 40nM for the 15, 20 or 34 nucleotide targets, respectively. Briefly, 2µL target was mixed with ∼10 000 microbeads in 10µL hybridization buffer (10× SSC, 0.1% SDS). The volume of each reaction was topped up to 20µL using water. The solution was incubated for 25 minutes at 37⁰C. Microbeads were magnetically washed using 200µL wash buffer. Washing was repeated twice before microbeads were resuspended in 250µL of PBS-T. Microbeads were transferred to flow tubes and individual microbead fluorescence was measured using a LSR II flow cytometer (Becton, Dickson and company, Franklin Lakes, USA) or LSR Fortessa (Becton, Dickson and company).

### Clinical samples

We obtained 64 patient plasma samples from University Health Network (Toronto, Canada). Samples were collected, stored and analyzed in accordance with our protocol (# 13-6974.13) approved by the University Health Network Research Ethics Board. We obtained second group of 53 deidentified hepatitis C virus, hepatitis B virus and COVID-19 samples. All patients provided written informed consent for the storage and use of their specimens for research. Samples were previously determined hepatitis C virus positive using the Abbott Realtime HCV assay (Abbott, Chicago, USA).

### Hepatitis C virus sequencing

Nucleic acid was extracted from each sample using QIAmp Viral RNA mini kits (Qiagen, Hilden, Germany) in accordance with manufacturer protocols. Extracted RNA was eluted into 45µL of water. A Nanodrop was used to determine the quantity and purity of the extracted RNA. RNA was frozen at −80⁰C for later use. RNA was reverse transcribed into cDNA using the SuperScript^TM^ VILO^TM^ cDNA synthesis kit (Invitrogen) in accordance with manufacturer protocols. cDNA was amplified using the Taq DNA polymerase with ThermoPol®-Buffer (New England Biolabs, Ipswich, USA). Briefly, 2.5µL 10× ThermoPol® reaction buffer, 0.2µL 25mM dNTPs, 0.5µL 10 µM forward primer (Table S3), 0.5µL 10 µM reverse primer, 0.125µL Taq DNA polymerase, 18.67µL nuclease-free water and 2.5µL template cDNA were combined in PCR strips. Each strip was spun down briefly and transferred to a preheated thermocycler. cDNA was denatured at 95⁰C for 30s followed by 35 cycles of 95⁰C for 20s, 54⁰C (core region) or 61.2⁰C (NS5B) for 50s and 68⁰C for 30s. PCR products were incubated at 68⁰C for 5 minutes then stored at 4⁰C.

Amplicons were purified using Ampure XP magnetic beads (Beckman Coulter, Brea, USA) and quantified using the Quant-iT^TM^ PicoGreen assay kit (Thermo Fisher). The NebNext® Ultra^TM^ II DNA library prep kit (NEB) for Illumina was used for sequencing library preparation following manufacturer’s instructions. Libraries were dual-barcoded with the NEBNext® Multiplex Oligos for Illumina® (Dual Index Primers Set 1). The libraries were pooled in equal amounts, after which the final pool was denatured and diluted to a sequencing concentration of 14pM. We sequenced samples using an Illumina MiSeq Nano kit with 2 × 250 bp paired end reads. Sequencing was demultiplexed on the MiSeq and the raw sequence data was processed with trimmomatic ^29^ to remove adapters and low-quality sequences. The pre-processed reads were analyzed for mutations using the virus variant detection pipeline virvarseq. ^30^ The reference sequences used for the virvarseq can be found in GenBank (GenBank accession numbers: NC_038882.1, NC_009823.1, NC_009824.1, NC_009825.1, NC_009826.1 and NC_009827.1). The coding frame of the sequences were determined after translation on all 6 frames using the expasy online tool. ^31^ The assembled sequences for each sample were compared to identify the greatest read depth. The sequences with the greatest read depths were aligned to identify 20 nucleotide regions with mutations and high sequence coverage. The alignment for part of the hepatitis C virus core gene can be seen in Figure S7. 20-nucleotide regions starting at position 574 and 8572 relative to reference genome H77 were chosen for the core and NS5B probes, respectively.

### Clinical virus detection

Hepatitis C virus and SARS-CoV-2 virus was extracted and reverse transcribed as described above. Hepatitis B virus was extracted from fine needle aspirates or whole blood using the DNeasy blood and tissue kit (Qiagen). cDNA from clinical samples (Table S2, S3) was PCR amplified using Taq DNA polymerase with ThermoPol®-Buffer (New England Biolabs). Briefly, 2.5µL 10× ThermoPol® reaction buffer, 0.2µL 25mM dNTPs, 0.5µL 10µM Cy5-labelled forward primer, 0.5µL 10µM reverse primer, 0.125µL Taq DNA polymerase, 18.17µL nuclease-free water and 3µL template cDNA were combined in PCR strips. Primers can be found in Table S4. Each strip was spun down briefly and transferred to a preheated thermocycler. cDNA was denatured at 95⁰C for 30s followed by 40 cycles of 95⁰C for 20s, 52⁰C for 40s and 68⁰C for 30s. PCR products were incubated at 68⁰C for 5 minutes and temporarily stored at 4⁰C. Quantum dot barcodes were synthesized and conjugated with amine-terminated DNA probes using EDC chemistry. PCR products were heat denatured for 5 minutes at 95⁰C. The 4µL of denatured DNA products were incubated with quantum dot barcodes (∼10 000 barcodes) in 16µL of hybridization buffer (10× SSC, 0.1% SDS). The total solution volume was 20µL. Tubes containing the solution were vortexed then were incubated for 25 minutes at 37⁰C on a rotator. Tubes were transferred to magnetic racks for 10 minutes and washed (0.5× SSC, 0.1% SDS) twice. Washed barcodes were resuspended in 250µL of PBS-T (1× PBS, 0.05% Tween-20) and transferred to flow tubes. Each sample was run in duplicate. Microbead fluorescence was measured using a LSR II flow cytometer.

### Effective match strategy

We increased clinical sensitivity using the effective match strategy. The effective match strategy improves the clinical performance of a given diagnostic probe by building that single probe into a panel of probes. The panel detects different variants of the same target that arise through viral mutation. The effective match strategy operates using the following assumptions: 1) Targets with less than 20% mismatches to a probe (referred to as effective matches) can be detected by that probe. Targets with 20% mismatches or more are undetected targets. 2) Once a probe is added to the panel, targets are removed from the pool of undetected targets if they have less than 20% mismatches to that probe.

The effective match strategy maximized the number of effective matches by selecting probes to add to the multiplex probe panel. The steps in the effective match strategy were as follows. The first step was to identify the sequence of the starting probe. We then aligned the probe sequence to target sequences obtained from a sequence database. Sequence databases can be private or public (*i.e.* NCBI Nucleotide). We identified mismatches between the probe and the diverse viral targets in the sequence database and calculated the number of undetected targets. We designed a library of probes capable of detecting the undetected targets. To select the second probe for the multiplex, we identified mismatches between candidate probes and the undetected targets. We then selected the probe that had <20% mismatches to the largest number of undetected targets. If there was a tie between probe candidates, we chose the probe with the most positive self-dimer ΔG. Self-dimer ΔG was predicted using the OligoAnalyzer (Integrated DNA Technologies). The process of identifying and quantifying mismatches between candidate probe and yet undetected target sequences was repeated to add additional probes to the multiplex panel.

### Analysis

The number of space between and position of mismatches in 15-mer targets was quantified using Microsoft Excel 2010 (Microsoft, Seattle, USA). Relative binding data was grouped so that position, number and space between mismatches was equally represented. A sigmoidal model and a quadratic model were fit to the mismatches number and position data, respectively. A Kruskal-Wallis test was used to compare groups of samples with zero, one or two spaces between mismatches. Non-linear regression was done using GraphPad Prism V7 (GraphPad Software, La Jolla, USA) and JMP Pro for Windows V14 (SAS Institute Inc., Cary, USA).

Flow cytometry data was analyzed on FlowJo V10 (FlowJo LLC, Ashland, USA) by gating the microbead population using FSC-A vs. SSC-A. Signal intensity was measured using the median B530/30 (fluorescein) or R660/20 (Cy5) filters. Each run for synthetic targets contained a standard curve of the perfect match target. Background signal was subtracted from all targets. Signal from non-perfect match targets was normalized to the maximum signal from the standard curve. Synthetic targets with <10% signal relative to the perfect match target for the same probe were considered negative. The limit of detection for clinical targets was calculated by averaging the background fluorescence of the negative samples and adding three standard deviations. Clinical samples with fluorescence signal less than the limit of detection were considered negative. Clinical sensitivity was calculated for all clinical samples and for targets with a given quantity of mutations. Signal intensity and clinical sensitivity relative to mismatch characteristics were plotted using GraphPad Prism V7. A Chi-square approximation for the Kruskal-Wallis test was used to evaluate whether there was a difference in sensitivity when using one to five probes. Differences between specific groups was determined using the Tukey-Kramer multiple comparison of the means.

## SUPPORTING INFORMATION

Additional experimental data, DNA sequences and clinical specimen information.

## Author contributions

H.N.K contributed to conceptualization, analysis, investigation, methodology, project administration, writing-original draft, and writing-review & editing. A.A.L. contributed to analysis, investigation and writing-review & editing. M.A.Z. contributed to investigation and writing-review & editing. J.J.F. contributed to resources and writing-review & editing. W.C.W.C. contributed to conceptualization, funding acquisition, resources, supervision, writing-original draft and writing-review & editing.

## ACKNOWLEDGEMENTS

W.C.W.C. acknowledges the Temerty Faculty of Medicine, Canadian Research Chairs Program Grant 950-223824 and Connaught ISI (313897) for funding support. H.N.K. would like to thank Paul Cadario, Mr. & Mrs. Ruggles, the Vanier Canadian Graduate Scholarship from CIHR, the McLaughlin Institute and the University of Toronto MD/PhD program. We thank the University of Toronto Centre for the Analysis of Genome Evolution and Function (CAGEF) for their genome sequencing and analysis services. We also thank Ayden Malekjahani for the fruitful discussions.

## SUPPLEMENTARY SECTION

**Figure S1.**
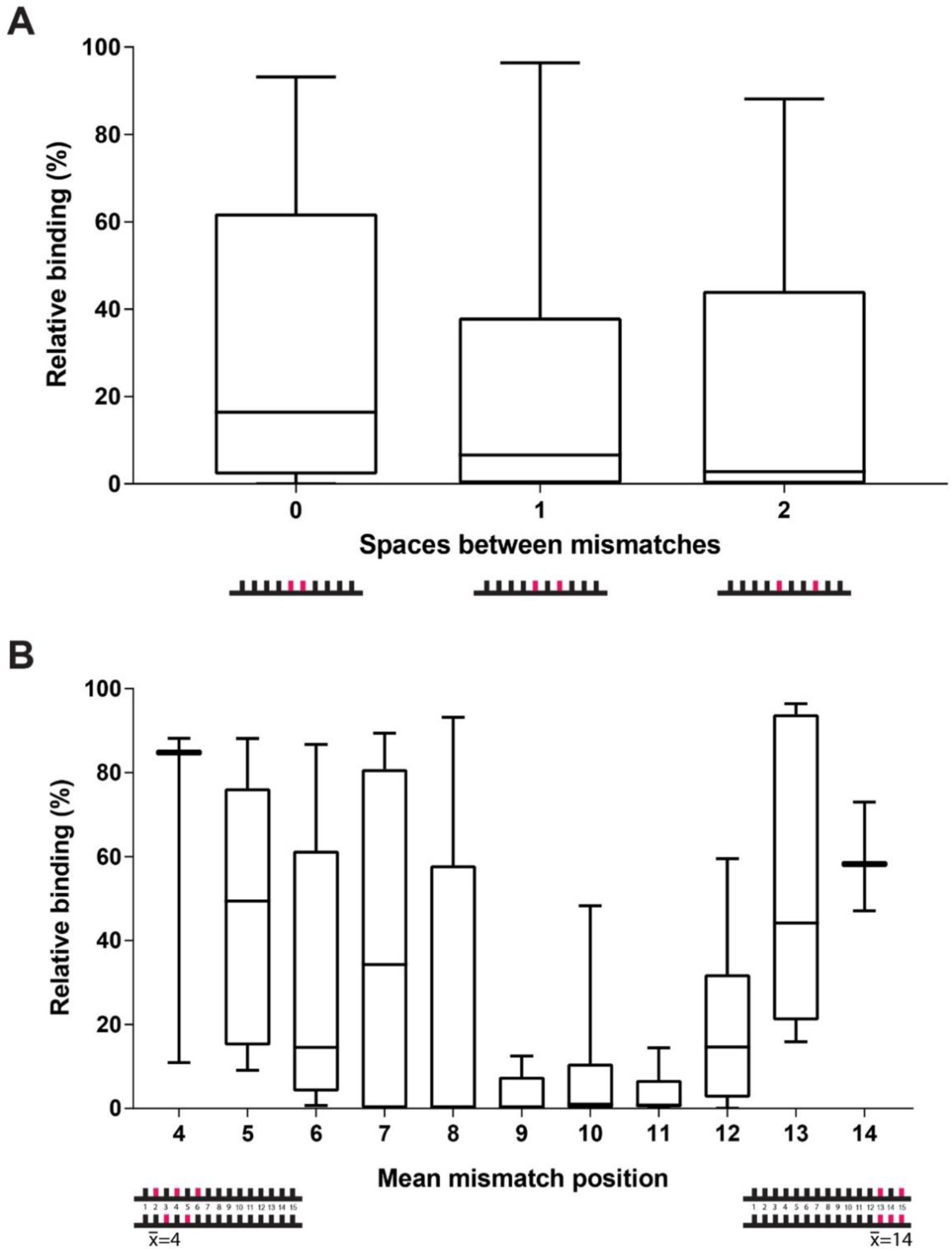
Effect of mismatch spacing and position on relative binding using microbead-conjugated probes and fluorophore-labelled targets. A) Box plot depicting relative binding versus mismatch spacing. Mismatches were adjacent (0 space), separated by 1 matching base pair or separated by 2 matching base pairs. B) Box plot depicting relative binding versus mean mismatch position. Mean position was determined by identifying the mismatch position of all mismatches within a target, adding the position together and dividing by the total number of mismatches in the target. Measurements were relative to signal from the probe and its perfectly complementary target. Box plots depict the interquartile range.

**Figure S2.**
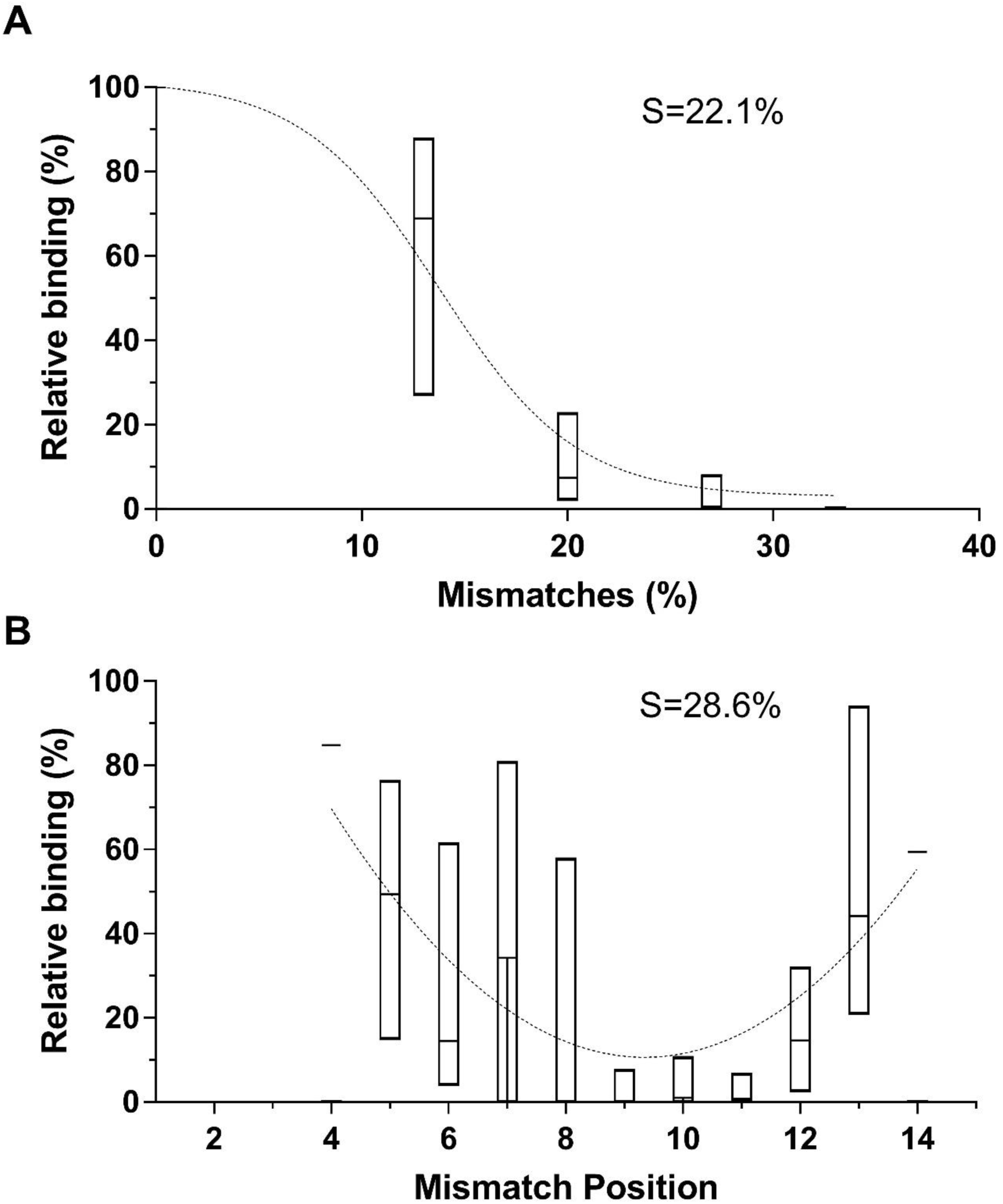
Modeling the effect of mismatch quantity and position on relative binding using microbead-conjugated probes and 15-mer targets. A) Sigmoidal curve modelling the effect of mismatch quantity on relative binding. B) Quadratic model of the effect of mismatch position on relative binding. Box plots depict the interquartile range. S is the standard error of the regression.

**Figure S3.**
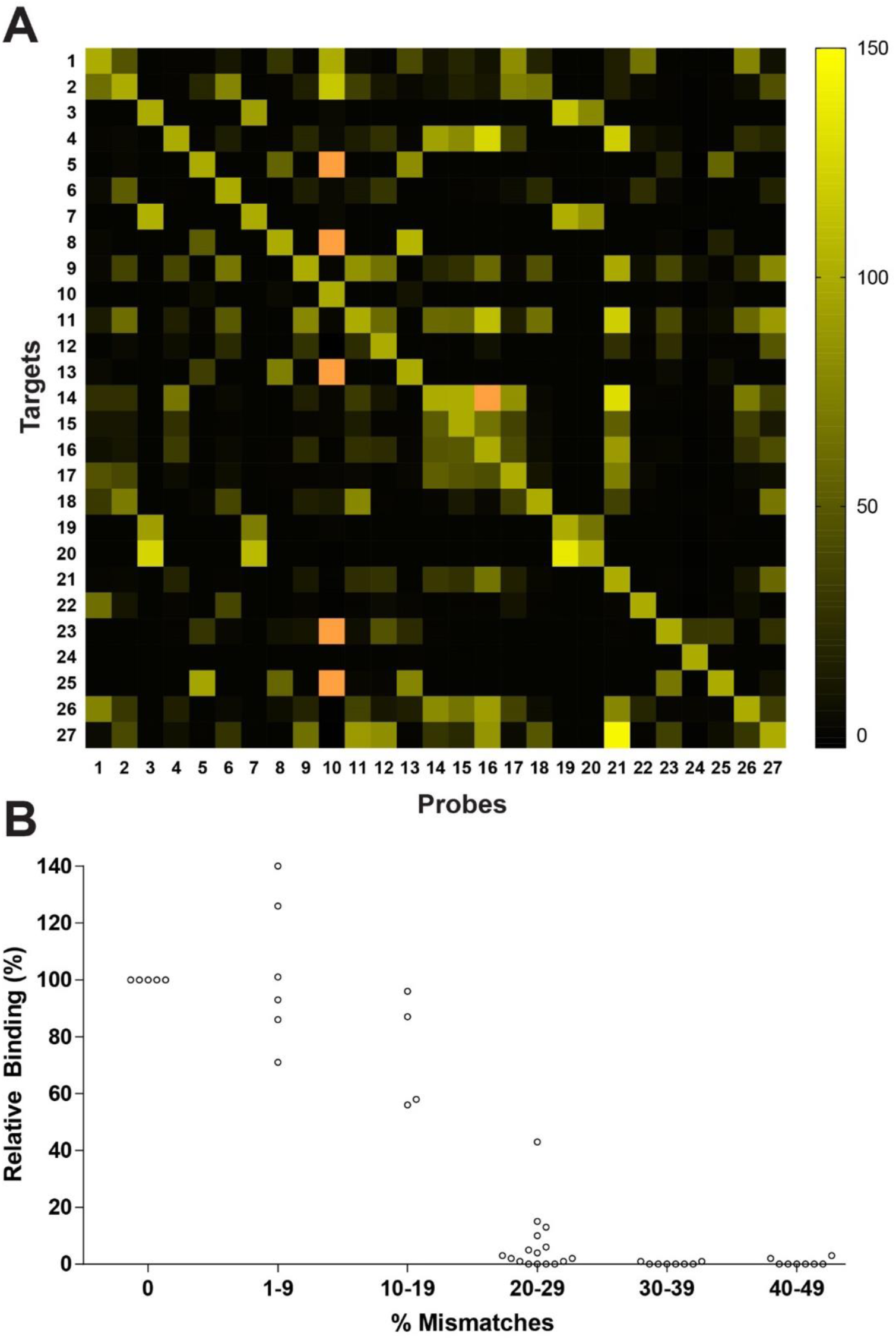
Measuring probe-target binding for 20- and 34-nucleotide microbead probes. A) The normalized fluorescence from 729 unique 20-mer probe-target combinations. The diagonal shows perfectly complementary probe-target pairs (100% signal). Orange indicates signal greater than 150% relative to the perfectly complementary sequence for that probe. B) Relative binding for 34-mer probe-target pairs with increasing numbers of mismatches. Fluorescence intensity was normalized to signal from the probe and its perfectly complementary target.

**Figure S4.**
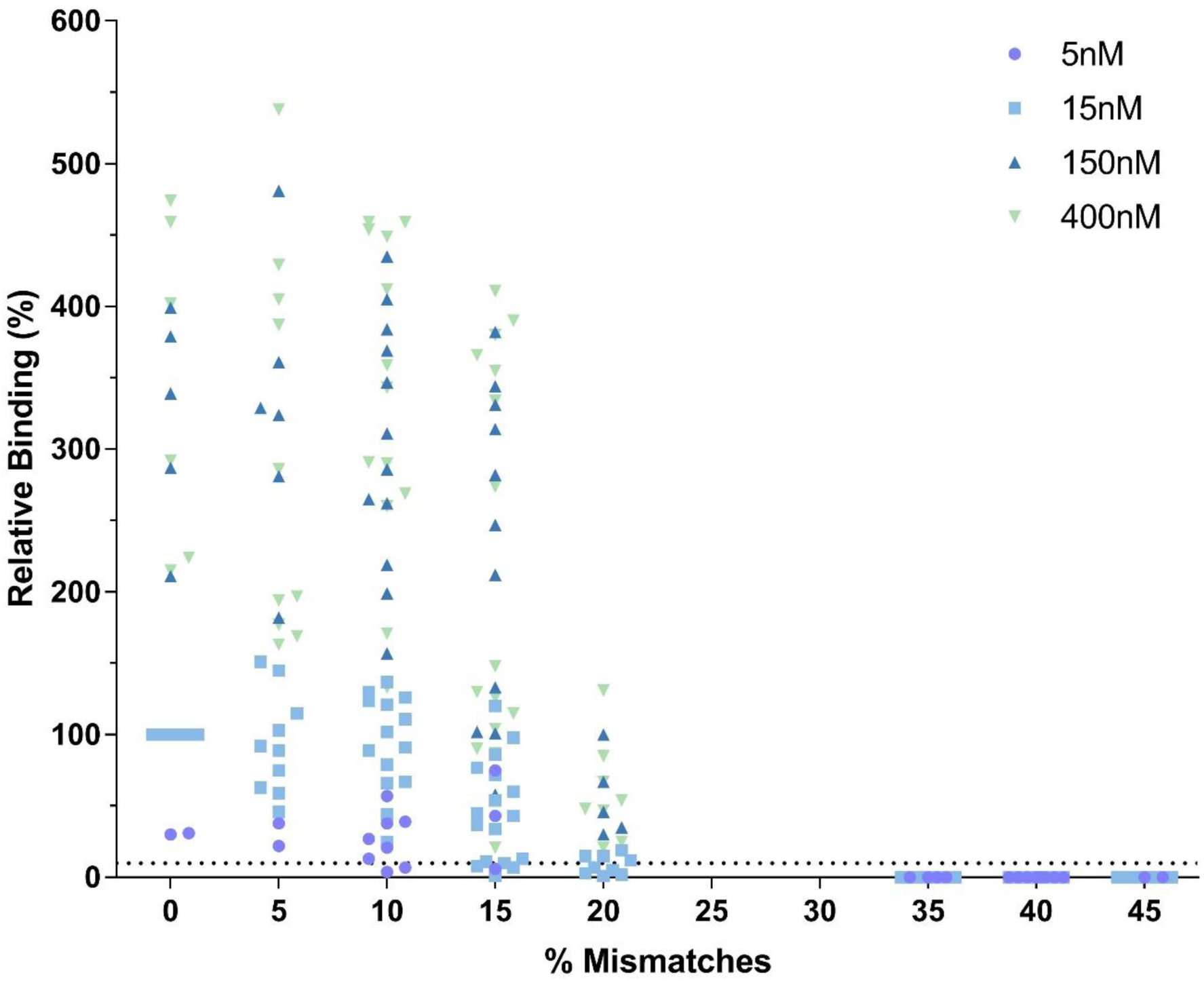
Measuring the effect probe-target mismatches on relative binding using targets at different concentrations. Relative binding decreased as the number of mutations increased. This was seen for all concentrations. Target concentrations of 5nM (purple circle) and 15nM (blue square) are within the linear range while 150nM (blue triangle) and 400nM (green triangle) are above the linear range. Binding is relative to 15nM perfectly complementary probe-target pairs (100% signal).

**Figure S5.**
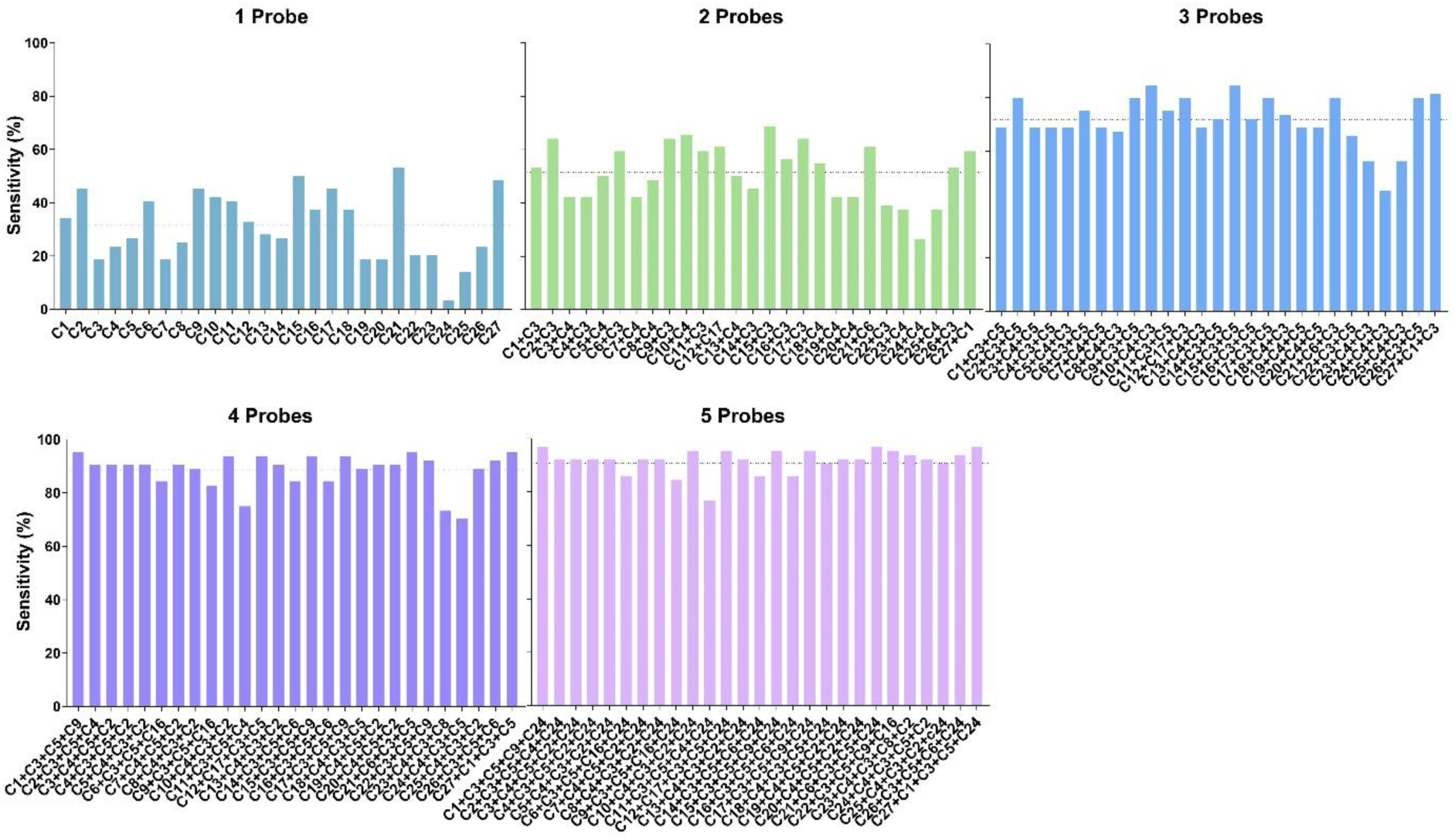
The clinical sensitivity achieved by combining probes to maximally increase the proportion of effective matches. We determined the effect of probe combinations on the clinical sensitivity. The probe sets are designed to detect multiple targets. For example, a 5-probe panel consisting of different emitting microbeads conjugated with sequences C1, C3, C5, C9 and C24 would perfectly bind to targets C1, C3, C5, C9 and C24. The detection of the target sequence enables us to determine the clinical sensitivity (equation 2). The specific probes are listed in Table S1. The dotted line represents the average across all combinations in that set.

**Figure S6.**
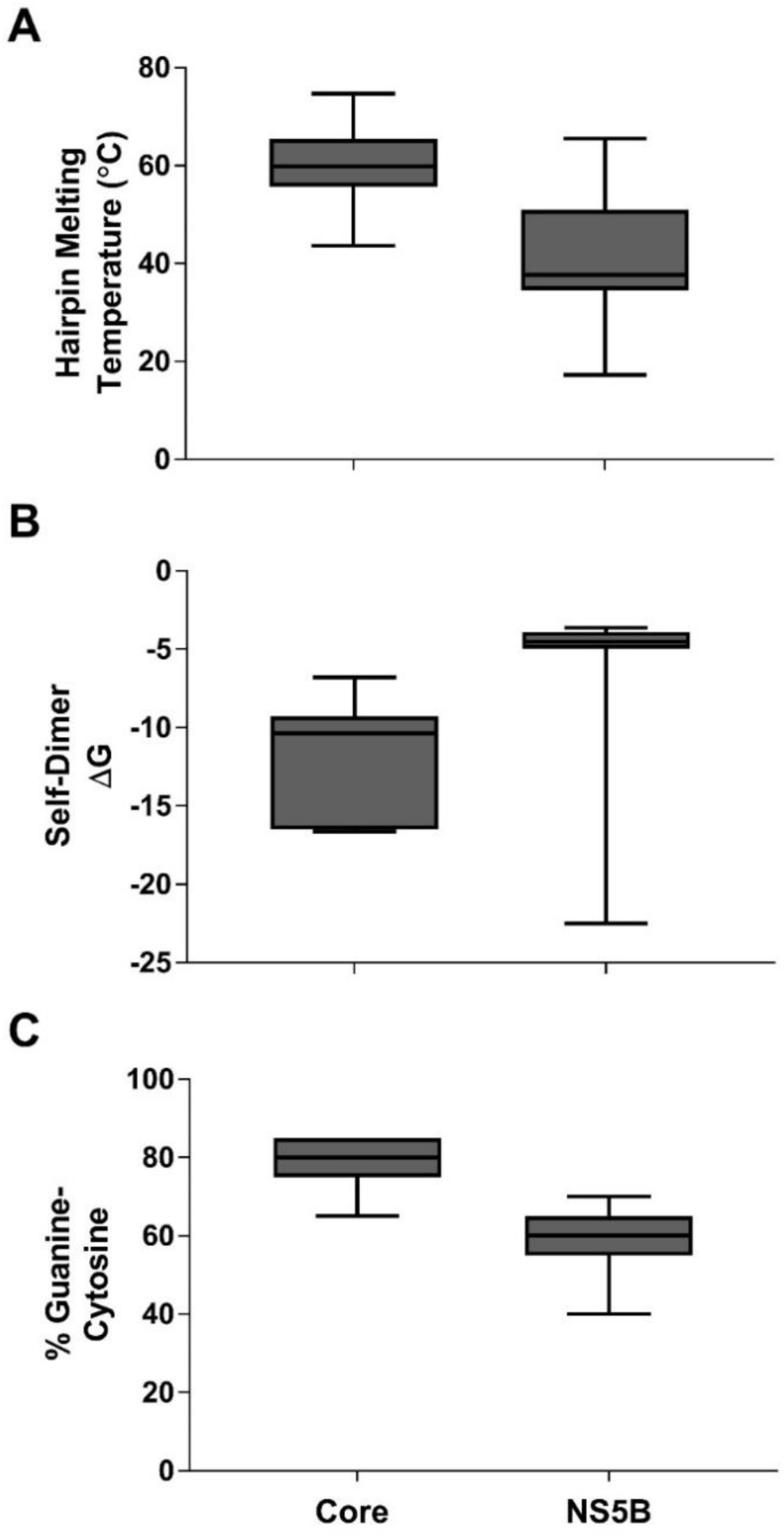
Structural characteristics of core and NS5B probe sets. Box plots of the A) hairpin melting temperature B) self-dimer free energy and C) percent guanine/cytosine (%GC) content of all core and NS5B probes. Core probes had more stable hairpins and were more likely to form self-dimers. Box plots depict the interquartile range.

**Figure S7.**
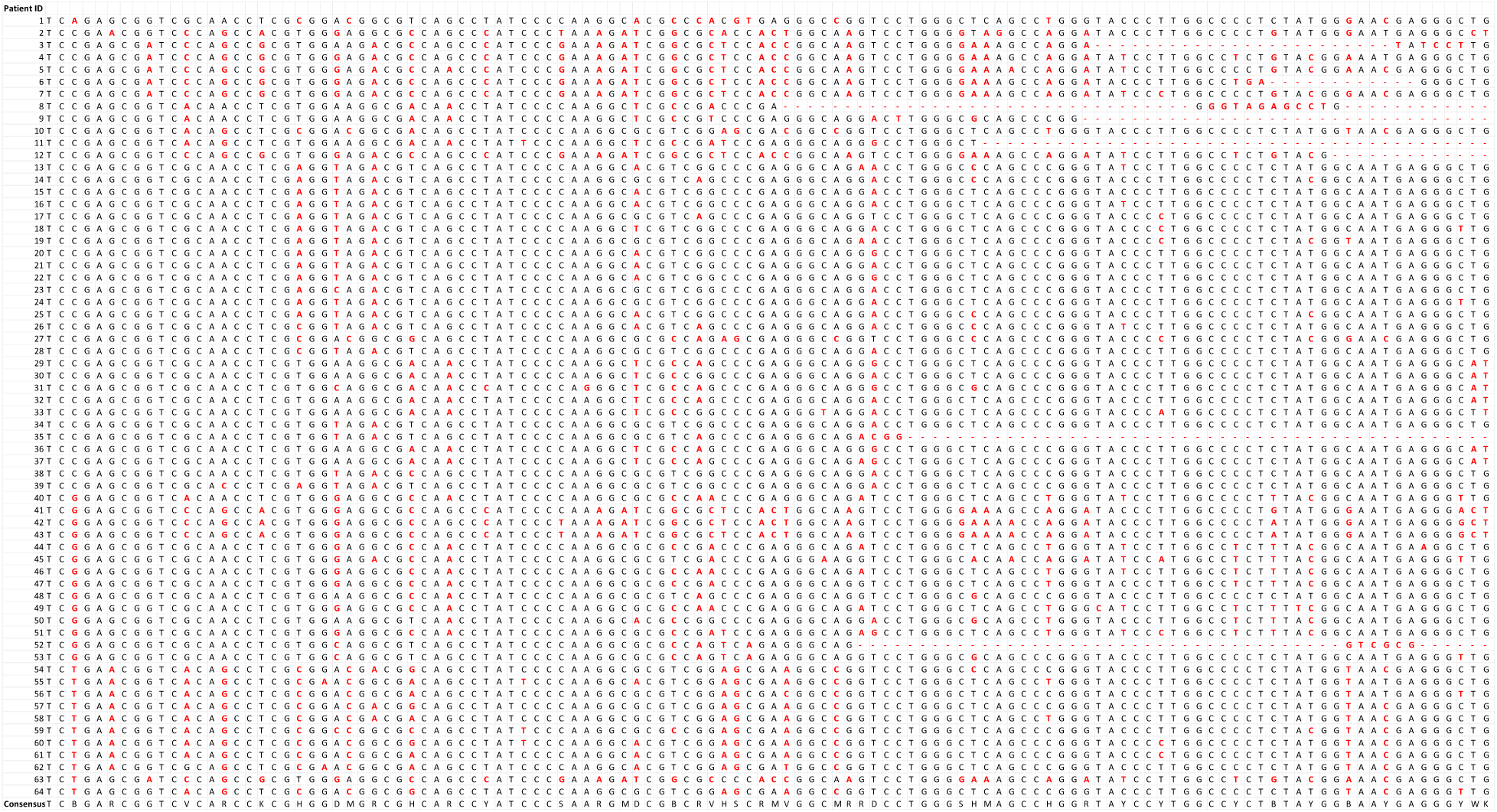
Alignment of 64 patient sequences for the hepatitis C virus core region. Each row represents a different patient sequence. The most common nucleotide in each position is in black and other nucleotides or gaps (-) are in red. The consensus sequence is written in the bottom row. RNA was extracted, reverse transcribed and PCR amplified prior to sequencing.

**Table S1.**
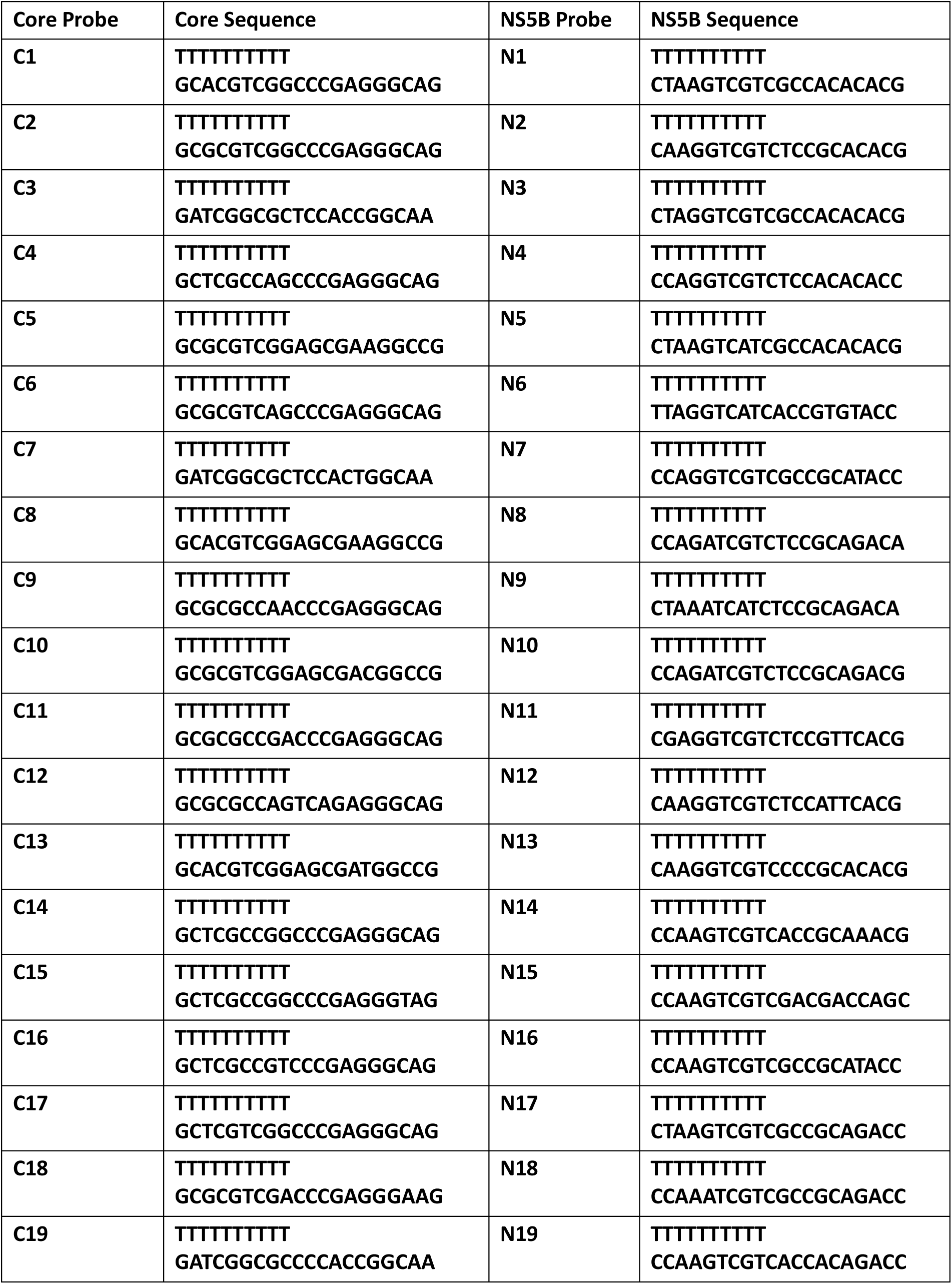
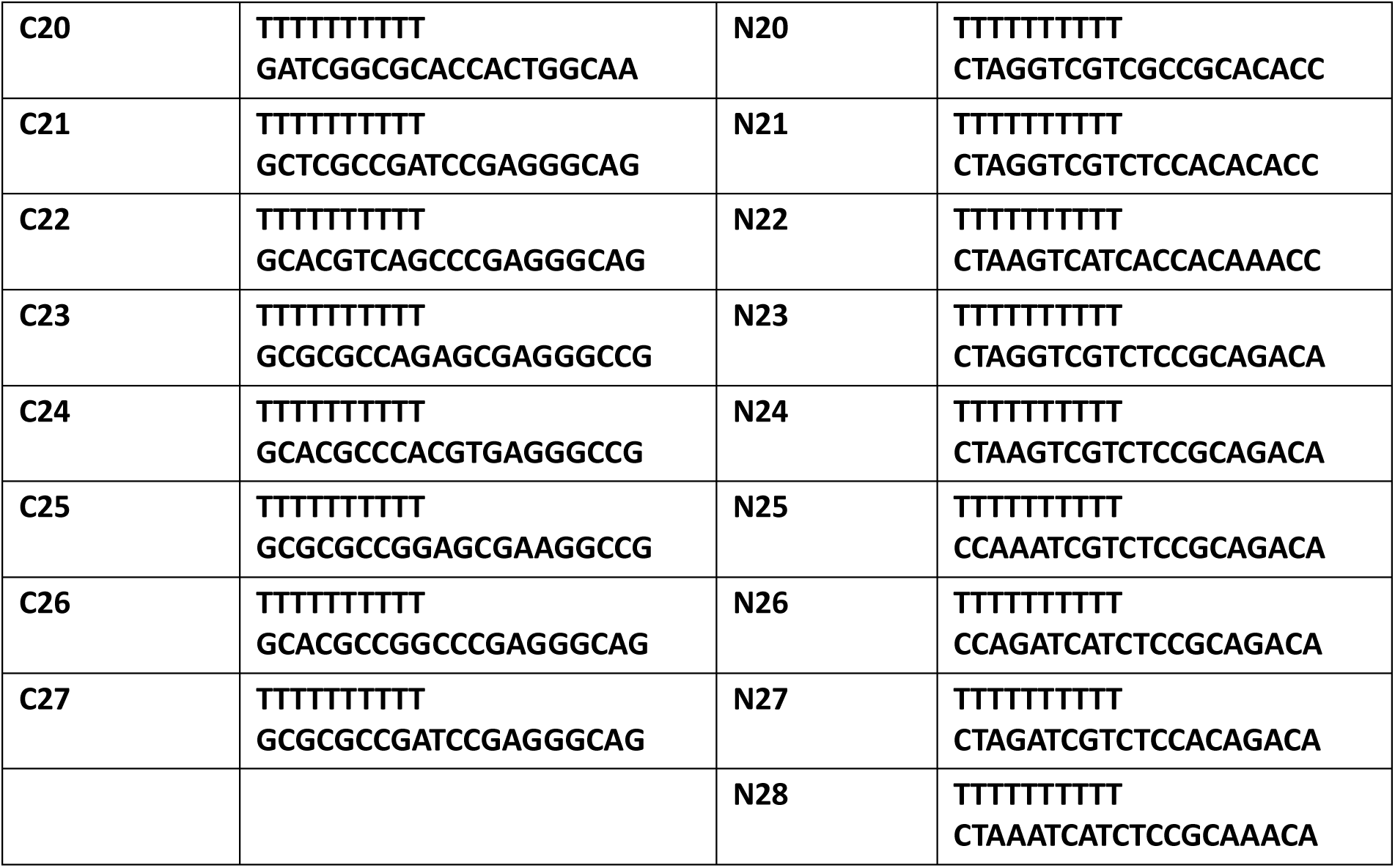
Nucleotide sequences for capture probes targeting the hepatitis C virus core and NS5B regions.

**Table S2.** Hepatitis C virus clinical samples tested using the N2 and N17 probes. Viral load as determined using the Abbot Realtime HCV assay.

| Sample ID | Genotype | Viral load (IU/mL) |
| --- | --- | --- |
| V1 | 1a | 4.59E+06 |
| V2 | 1a | 5.06E+06 |
| V3 | 1a | 1.69E+02 |
| V4 | 1a | 7.43E+05 |
| V5 | 1a | 9.47E+02 |
| V6 | 1a | 1.17E+03 |
| V7 | 1a | 1.67E+02 |
| V8 | 1a | 1.31E+06 |
| V9 | 1a | 1.05E+06 |
| V10 | 1a | 1.75E+07 |
| V11 | 1a | 1.69E+04 |
| V12 | 1a | 1.69E+04 |
| V13 | 1a | 1.00E+08 |
| V14 | 1a | 1.72E+06 |
| V15 | 1a | 6.19E+05 |
| V16 | 1a | 1.90E+05 |
| V17 | 1a | 3.00E+06 |
| V18 | 1a | 2.52E+04 |
| V19 | 1b | 1.38E+06 |
| V20 | 2 | 4.36E+02 |
| V21 | 2 | 3.25E+06 |
| V22 | 2 | 1.17E+06 |
| V23 | 3 | 2.02E+02 |
| V24 | 3 | 5.27E+02 |
| V25 | 3 | 2.00E+01 |
| V26 | 3 | 1.18E+06 |
| V27 | 3 | 1.54E+07 |
| V28 | 3 | 2.40E+03 |
| V29 | 3 | 5.48E+06 |
| V30 | 3 | 5.48E+06 |

**Table S3.** Hepatitis B virus and SARS-CoV-2 clinical samples tested using the N2 and N17 probes. Viral loads for Hepatitis B virus were between 10^7^-10^8^ copies/mL and for SARS-CoV-2 were between 10^2^-10^10^copies/mL.

| Sample ID | Virus | Detected by N2 | Detected by N17 |
| --- | --- | --- | --- |
| H1 | HBV | **Negative** | Negative |
| H2 | HBV | Negative | Negative |
| H3 | HBV | Negative | Negative |
| H4 | HBV | Negative | Negative |
| H5 | HBV | Negative | Negative |
| H6 | HBV | Negative | **Positive** |
| H7 | HBV | Negative | Negative |
| H8 | HBV | Negative | Negative |
| H9 | HBV | Negative | Negative |
| H10 | HBV | Negative | Negative |
| H11 | HBV | Negative | **Positive** |
| S12 | SARS-CoV-2 | Negative | Negative |
| S13 | SARS-CoV-2 | Negative | Negative |
| S14 | SARS-CoV-2 | Negative | Negative |
| S15 | SARS-CoV-2 | Negative | Negative |
| S16 | SARS-CoV-2 | Negative | Negative |
| S17 | SARS-CoV-2 | Negative | Negative |
| S18 | SARS-CoV-2 | Negative | Negative |
| S19 | SARS-CoV-2 | Negative | Negative |
| S20 | SARS-CoV-2 | Negative | Negative |
| S21 | SARS-CoV-2 | Negative | Negative |
| S22 | SARS-CoV-2 | Negative | Negative |
| S23 | SARS-CoV-2 | Negative | Negative |

**Table S4.**
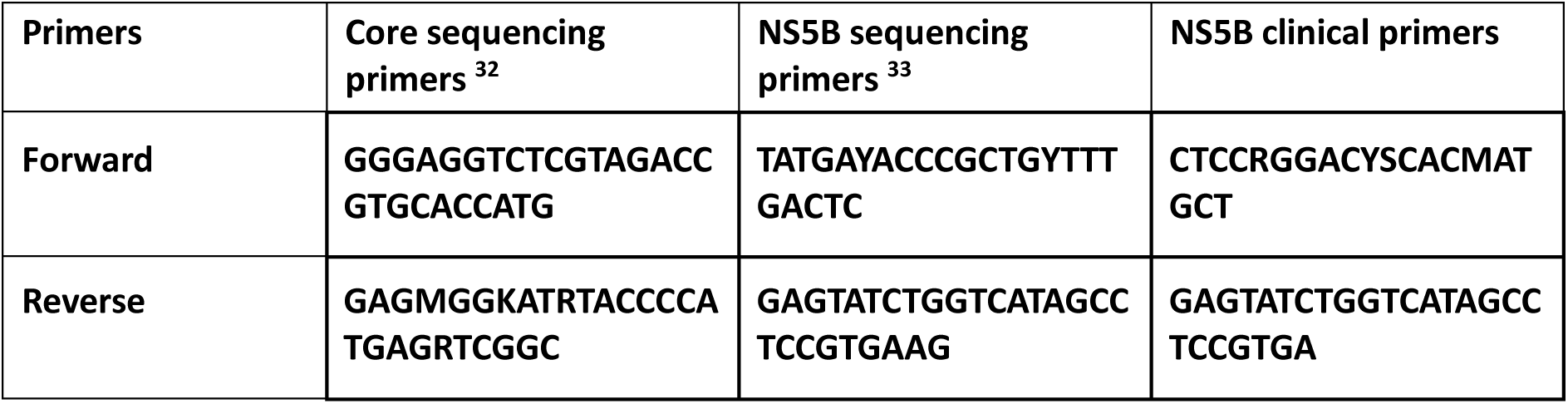
Primer sequences used for amplifying the core and NS5B regions of the hepatitis C virus.

